# The ants (Hymenoptera: Formicidae) of Iceland

**DOI:** 10.64898/2026.11.05.764515

**Authors:** Marco Mancini, Andreas Guðmundsson Gähwiller, James K. Wetterer, Arnar Pálsson

## Abstract

The native terrestrial biota of Iceland required long oceanic journeys to colonize the island. Human colonization of Iceland, however, has allowed many less vagile species, such as ants, to reach the island. We compiled published, unpublished, and new records of ants from Iceland and evaluated the probable origin and current status of each species. We analyzed 565 records compiled by the Icelandic Institute of Natural History (1974-2020) and 84 new records (2019-2023) representing 25 species. Only five species were found living outdoors: *Hypoponera ergatandria* (507x indoors and 9x outdoors, mostly in geothermal areas), *Lasius niger* (24x indoors and 12x outdoors away from heated buildings), *Hypoponera eduardi* (5x only in geothermal areas), *Lasius platythorax* (1x near a small greenhouse and 1x in imported plant material), and *Myrmica rubra* (1x in an urban garden). Three ant species were exclusively found inhabiting heated buildings: *Monomorium pharaonis* (41x), *Tapinoma melanocephalum* (10x), and *Monomorium* nr. *pharaonis* (3x). Records of the remaining 17 species all appear to be based on temporary infestations, found on recently imported plant material. Previous records from Iceland of *Hypoponera punctatissima* appear to be based on misidentifications. Although it is possible that some ants were present before human arrival, all 25 ant species may have arrived accompanying humans. It remains to be seen whether any of these ant species are having or will have a significant impact on the native Icelandic biota. The arrival of *Myrmica rubra*in Iceland is particularly worrisome, given the impact this species is having in other high latitude areas it has invaded.

## Introduction

Iceland is a volcanic island located in the North Atlantic on the Mid-Atlantic Ridge (DOWNES 1988; TORSVIK & al. 2001; LUNDIN & DORÉ 2002). The nearest landmasses are Greenland (∼280 km), the Faroe Islands (∼420 km), Scotland (∼800 km), and Norway (∼970 km). During the Last Glacial Maximum, 20-18kya, nearly all of Iceland was buried in ice. Most of the native terrestrial biota colonized Iceland after the glaciers began to retreat, feats that required long oceanic journeys. Human colonization of Iceland ∼1.2kya, however, opened the opportunity for many less vagile species to reach the island. In fact, most insect species now found in Iceland and other North Atlantic islands appear to have arrived in the past 1.2ky (DUGMORE & al. 2005). In the past 50y, increases in travel and imports has resulted in great increases in the arrival of additional non-native species (MINISTRY FOR THE ENVIRONMENT & THE ICELANDIC INSTITUTE OF NATURAL HISTORY 2001). In the present study, we compiled published, unpublished, and new records of ants from Iceland and evaluated the probable origin and current status of each species.

Of the 14.438 extant ant species known (ANTCAT 2026), over 200 have established populations outside their native range (MCGLYNN 1999). Several of these species are highly invasive, with significant ecological, environmental, and economic impacts (HOLWAY & al. 2002). Historically, most invasive ant species originated in tropical and subtropical areas. However, ant invaders are also now spreading from temperate ecoregions (SUAREZ & al. 2010). This, and global warming, may expose Iceland and other North Atlantic islands to ant invasions. For example, six alien ant species were recently detected in the Faroe Islands, three of which are native to temperate ecoregions (HAMMER & JENSEN 2021). In many remote oceanic islands, the native fauna is less diverse and abundant compared to neighboring mainland areas (PAULAY 1994). When ants invade such islands, they can prey on naïve endemic fauna and devastate local ecosystems (PASSERA 1994), as seen on native invertebrates in volcanic areas of Hawaii (MEDEIROS & al. 1986; WETTERER 1998) and on Christmas Island (BAUMGARTNER & RYAN 2020).

Published records of ants in Iceland are very limited. The first mention dates to PETERSEN (1956), who recorded only *Ponera punctatissima* (= *H. punctatissima*). ÓLAFSSON & RICHTER (1985) reported 75 records *H. punctatissima* in the city of Reykjavík between 1974 and 1984, mainly in basements and ground floors in the old town. BUCKLAND & al. (1992) reported specimens of *H. punctatissima* and *Leptothorax* sp. in excavated layers dated to the 17^th^ or 18^th^ Century in Reykholt (Borgarfjörður). In addition, MANCINI & al. (2022) recently reported a *Myrmica rubra* colony in one garden in Reykjavík.

SEIFERT (2013, 2018) found that some specimens identified as *H. punctatissima* were actually a separate species, *H. ergatandria*, with the latter showing greater worldwide dispersal. This taxonomic revision suggests a need to re-evaluate Icelandic records of ants previously identified as *H. punctatissima*.

## Materials and methods

Records of ants in Iceland came from two sources: historical records from the Icelandic Institute of Natural History (hereafter IINH) and newly collected University of Iceland (hereafter UI) specimens. The staff of IINH are regularly contacted by pest management professionals and citizens seeking pest species identifications or reporting insect findings. These records are processed by museum staff, sometimes identified to species or genus level with the help of experts. In January 2020, we reviewed all ant records in this database, which includes species identification, collection site (indoor/outdoor and postal codes), date and year, number of specimens collected, and general notes about the findings.

In addition, we contacted eight pest management professionals operating in Reykjavík and the neighboring townships in the summer of 2019, and asked if they had encountered ants. Seven reported that they rarely encountered them, mainly because their job focused on other animal infestations (e.g., mice, rats, silverfish, bed bugs). However, one agent (hereafter SSG), reported that he regularly handled ant cases, and allowed us to join him in his operations. For each ant-case we recorded: ant species, date, time, street address, GPS coordinates, and some general information regarding the finding (all records received a serial number). During the same period, we developed a website to raise public awareness (http://maurar.hi.is) under the auspices of the Institute of Life and Environmental Sciences at the University of Iceland, with information on the biology and distribution of the ants in Iceland) and linked social media accounts to provide information about ants and solicit the public to report ants.

### Specimen identification

Erling Ólafsson identified all the specimens collected by the IINH with the help of international experts, and voucher specimens are available in the Institute’s collections. Two of the specimens were only identified to genus, and two others to subfamily. Most of the new specimens (after 2019) were identified by Bernhard Seifert (Senckenberg Research Institute. Görliz, Germany). In addition, John T. Longino (University of Utah. Salt Lake City, USA) identified *Pheidole* sp. and John LaPolla (Towson University. Towson, USA) *Nylanderia* spp. specimens. Voucher specimens have been deposited in the UI’s collections. Of those, 71 specimens collected between 7 August 2019 and 14 December 2021 were sent to the Canadian Centre for DNA Barcoding (CCDB) in Guelph, Canada, for sequencing of the *COX1* gene. To compare the IINH and new datasets, we proceeded with the re-identification (performed by Seifert) of 16 historical records found in the IINH database, and originally identified as *H. punctatissima*. These records span from 30 September 1977 to 2 October 2018 and represent six locations in four different regions of the country.

### DNA sequence analyses

Records were summarized with table summaries and plots were created with R statistical environment, version 4.2.0 (R CORE TEAM 2022). Matches of the 69 DNA sequences were found in the National Center for Biotechnology Information (NCBI) using Blast (October 2024). One or more sequences that best matched the Icelandic samples, usually with a >95% similarity, were compiled in the Mesquite software, version 3.81 (MADDISON & MADDISON 2023), names were standardized, with species designations and sampling locations where applicable. Alignment was done with ClustalW (THOMPSON & al. 1994), within the MEGA software, version 11.0.13 (TAMURA & al. 2021). The jModelTest, version 2.1.10 (DARRIBA & at. 2012), was used to deduce the best-fitting evolutionary model for the data, in this case GTR+I+G. Finally, a phylogenetic analysis was conducted in MrBayes, version 3.2.7 (HUELSENBECK & RONQUIST 2001), with the previously determined parameters. The results from these analyses were visualized in the Dendroscope software, version 3.8.10 (HUSON & SCORNAVACCA 2012).

All sequences from the study are available in NCBI, (XXX-XXX), except for *Myrmica rubra* (NCBI, ON131872.1), that was previously submitted (MANCINI & al. 2022).

## Results

In total, we analyzed 649 Icelandic ant records, representing 25 ant species detected over the past six decades (Tab. 1). Historical records from the literature were set aside from our main dataset to avoid biasing the spatial and temporal trends. Specifically, we excluded PETERSEN’s (1956) record due to imprecise locality metadata, while the 73 records from ÓLAFSSON & RICHTER (1985) were left out of statistical testing to keep the focus strictly on modern records (1985–2023). Likewise, the two archaeological specimens (*H. punctatissima/ergatandria* and *Leptothorax* sp.) reported from 17^th^ or 18^th^ Century layers in Reykholt (BUCKLAND et al. 1992) require separate physical re-examination and were omitted from our frequency analyses.

**Tab. 1:** First occurrences of ants found in Iceland, as recorded by the Icelandic Institute of Natural History (IINH) and the University of Iceland (UI) collections. Species are listed chronologically by their earliest known records and locations (all towns or villages). The first year in which an ant species was documented in Iceland is highlighted in bold. In some cases, specimens were identified only to the genus level (e.g., *Pheidole* sp.) or subfamily level (e.g., *Myrmicinae*, species a).

| N. | Ant species | Collection |  | Location<br>of first finding |
|---|---|---|---|---|
|  |  | IINH | UI |  |
| 1 | *Hypoponera ergatandria* | **<1956** | 2019 | Borgarfjörður |
| 2 | *Colobopsis truncata* | **1977** | 2021 | Kópavogur |
| 3 | *Camponotus herculeanus* | **1980** |  | Þorlákshöfn |
| 4 | *Monomorium pharaonis* | **1980** | 2019 | Reykjavík |
| 5 | *Leptothorax acervorum* | **1983** |  | Reykjavík |
| 6 | *Hypoponera eduardi* | **1987** | 2020 | Geysir |
| 7 | *Camponotus vagus* | **1988** |  | Þorlákshöfn |
| 8 | Myrmicinae, species a | **1989** |  | Reykjavík |
| 9 | *Crematogaster scutellaris* | **1991** | 2020 | Reykjavík |
| 10 | *Lasius niger* | **1994** | 2019 | Reykjavík |
| 11 | *Lasius emarginatus* | **1996** |  | Reykjavík |
| 12 | *Pheidole* sp. | **1996** |  | Reykjavík |
| 13 | Myrmicinae, species b | **1999** |  | Kópavogur |
| 14 | *Trichomyrmex destructor* | **2000** |  | Seltjarnarnes |
| 15 | *Tetramorium caespitum* | **2009** |  | Reykjavík |
| 16 | *Tapinoma melanocephalum* | **2009** | 2021 | Reykjavík |
| 17 | *Camponotus* sp. | **2014** |  | Reykjavík |
| 18 | *Nylanderia bourbonica* |  | **2019** | Kópavogur |
| 19 | *Monomorium* nr. *pharaonis* |  | **2020** | Hafnarfjörður |
| 20 | *Pheidole bilimeki* |  | **2020** | Reykjavík |
| 21 | *Nylanderia jaegerskioeldi* |  | **2021** | Kópavogur |
| 22 | *Tetramorium bicarinatum* |  | **2021** | Garðabær |
| 23 | *Myrmica rubra* |  | **2021** | Reykjavík |
| 24 | *Lasius platythorax* |  | **2021** | Akranes |
| 25 | *Monomorium floricola* |  | **2022** | Reykjavík |

The IINH database included 565 ant records, corresponding to 17 species. All records but two were found in urban areas and mainly through personal communication of pest management professionals and citizens. Experts identified 13 taxa to species, two to genus (*Camponotus* sp. and *Pheidole* sp.) and two to subfamily (Myrmicinae, species a, and Myrmicinae, species b). We did not examine these IINH specimens. We were told that the four records not identified to the species level are distinct from the other identified species in the record. The dataset included records up to 5 February 2020, except for the records initially identified as *Hypoponera punctatissima*, which extended to 30 November 2020. Bernhard Seifert re-identified 16 records from IINH identified as *H. punctatissima* and found that 15 were *H. ergatandria* and one was *H. eduardi* (from the Southern Region). We therefore provisionally consider all previous records of *H. punctatissima*in Iceland to be misidentifications.

From 7 August 2019 to 6 March 2023, the UI sampling effort with a pest management professional (SSG) and the public yielded 84 records. Most records (49; 58.33%) came through the pest management professional, 16 (19.05%) records were reported by citizens and eight (9.52%) were accidental findings. The remaining 11 records (13.10%) represent ant surveys conducted in geothermal areas (10 records) and in a location close to an ant-site detected previously (one record). The UI records represent 15 species (Tab. 1) and were identified by international experts. Eight of the UI species were included in IINH records. In the UI records *H. ergatandria* was most common, while in the IINH it was *H. punctatissima*.

We sequenced a portion of the *COX1* gene in 69 specimens from 57 locations. In general, the sequencing data supported species designations and revealed low genetic diversity within each species in Iceland with none, one, or only a few variants separating specimens (Fig. 1). There were two notable exceptions. First, three of 12 *Monomorium* spp. sequences from Iceland formed two distinct clades with 90% sequence similarity. All specimens were initially identified based on external morphology as belonging to the same species, but this level of sequence divergence is normally associated with speciation. The clade with nine Icelandic sequences matched almost all other *M. pharaonis* records in the NCBI databases, but one sample from Germany (GU705833.1, identified as *M. pharaonis* from 1999) matched perfectly two of these three sequences. We suspect these three specimens were possibly misidentified as *M. pharaonis* and thus report them as unidentified *Monomorium* nr. *pharaonis.* Second, the Icelandic *H. eduardi* records perfectly matched sequences labelled *Hypoponera opacior* in GenBank, but only weakly *H. eduardi* sequences in the same database. We suspect both of these observations represent misidentified records in the database, because our records align well (99% identity) with multiple certified *H. eduardi* samples from the Mediterranean Area (MENCHETTI & al. 2026).

**Fig. 1:**
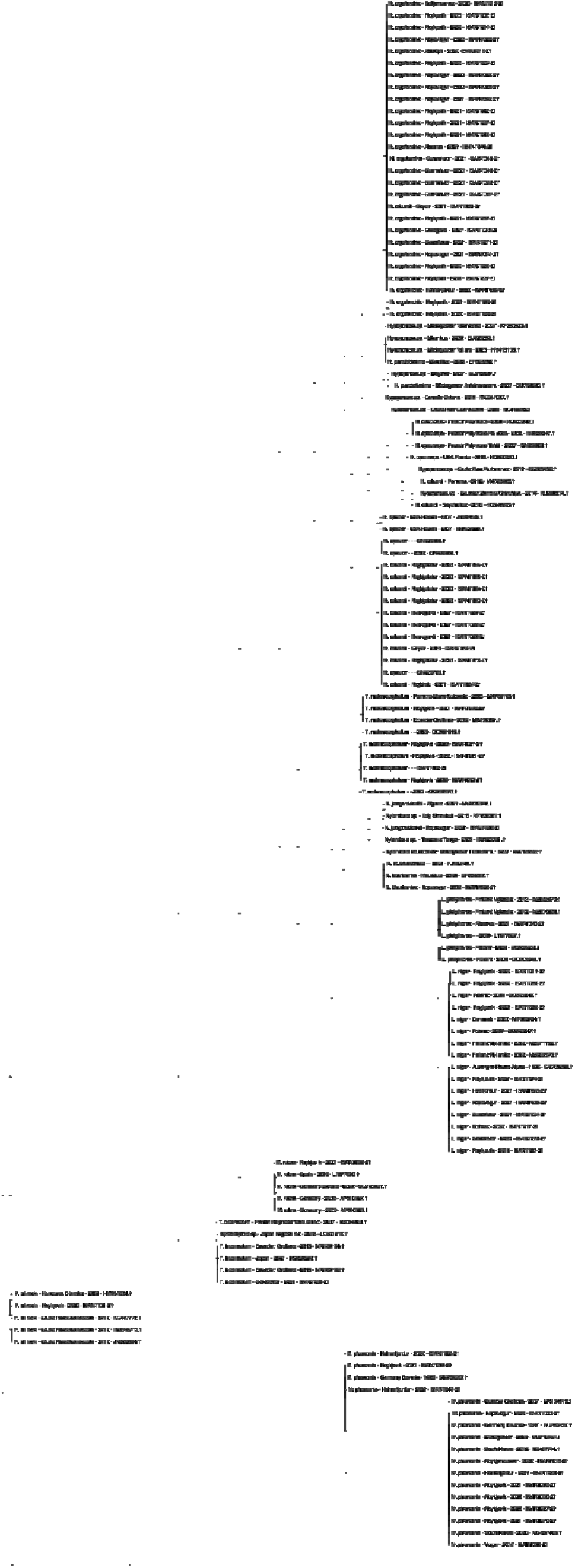
Unrooted phylogenetic tree of the 69 Icelandic records based on *COX1* gene, with one or more close hits from NCBI as representatives. The tree was generated with a jModelTest, best-fitting evolutionary model GTR+I+G. The phylogenetic analysis was conducted in MrBayes, version 3.2.7.

Overall, the genetic data show limited genetic variation within most of the species found in Iceland. Specifically, the *H. eduardi* showed no variation (low genetic diversity among sequenced samples), while *H. ergatandria* possessed two alleles (separated by a single base). The highest within-species variation was observed in *L. niger* records, along with some diversity found within *L. niger* and *T. melanocephalum*. A cautious interpretation of these results is that each species most likely colonized once, while repeated colonizations from the same (or a related) source cannot be disproven with these data. Most other cases represent colony fragments or colonies that have been exterminated on-site by pest management professionals and most likely represent casual alien species (PYŠEK & al. 2004).

### Species with known outdoor populations

*Hypoponera ergatandria* (FOREL, 1893) – IINH & UI. 516 records

Akranes (3, IINH; 1, UI), Akureyri (2, IINH; 1, UI), Blönduós (2, IINH), Dalvík (3, IINH), Flúðir (1, IINH), Garðabær (14, IINH; 2, UI), Grænidalur (1, UI), Gunnuhver (1, IINH; 1, UI), Hafnarfjörður (43, IINH; 1, UI), Hvammur (1, IINH), Hveragerði (3, IINH), Hvolsvöllur (1, IINH), Keflavík (5, IINH), Kleppjárnsreykir (1, UI), Kópavogur (34, IINH; 5, UI), Laugaland (1, IINH), Molsfellsbær (2, IINH; 1, UI), Ólafsfjörður (1, IINH), Reykhólar (1, UI), Reykholt, Borgarfjörður (1, IINH), Reykjavík (338, IINH; 20, UI), Sandgerði (1, IINH; 1, UI), Sauðarkrókur (2, UI), Selfoss (5, IINH), Seltjarnarnes (11, IINH; 2, UI), Varmahlíð (1, UI), Varmaland (1, IINH), Ytri-Njarðvík (1, IINH)

Native to the tropics and subtropics of the New World, *H. ergatandria* is considered a cosmopolitan tramp (SEIFERT 2013). Cryptic subterranean species common in greenhouses. Ergatomorphic gynes, winged gynes, and males have been observed. Polygynous, cryptic species with large numbers of winged gynes that emerge for dispersal. They possess a functioning stinger.

Occurrence in Iceland: Records of H. ergatandria account for ∼80% of total ant records (516 out of 649). Mostly in heated buildings in the urban environment, where swarming gynes are frequently detected attracted to light on windowsills. Very hard to eradicate colonies: to date, none of the *H. ergatandria* control treatments we participated in proved successful, and visible swarming only naturally ceased when gynes stopped emerging. Nine out of 516 cases were found outdoors, mostly in geothermal areas. Self-sustained populations have been documented for more than ten years.

*Lasius niger* (LINNAEUS, 1758) – IINH & UI. 36 records

Borgarnes (1, IINH), Garðabær (5, IINH; 2 UI), Húsafell (1, UI), Hvalfjörður (1, UI), Hveragerði (1, IINH), Kópavogur (1, IINH; 2, UI), Mosfellsbær (1, IINH), Reykjavík (11, IINH; 7, UI), Selfoss (1, UI), Vogar (1, IINH; 1 UI)

Eurosiberian species, widely distributed outdoors in temperate Europe. It is highly tolerant and adaptive to the urban environment and prefers to nest under flat stones. Strictly monogynous, it forms large colonies with up to 60.000 workers (SEIFERT 2018). Generalist, in terms of diet, tends aphids for honeydew and preys on small invertebrates.

Occurrence in Iceland: Large, mature colonies. In 12 of 36 cases, it was found outdoors (specifically in private gardens in Reykjavík, Garðabær, and Selfoss), usually with nests constructed under stones and workers foraging exclusively outdoors (on trees that host aphid populations). Treatment with sugar-based liquid baits proved 100% successful in controlling *L. niger* colonies.

*Hypoponera eduardi* (FOREL, 1894) – IINH & UI. 5 records

Geysir (1 IINH; 1 UI), Hveragerði (1), Reykholt, Bláskógabyggð (1), Reykjadalur (1)

Native to the Mediterranean area, where it is widely distributed (YARROW 1967; SEIFERT 2018). It is an alien and invasive species in several other countries around the world (COLLINGWOOD 1985; DON 2007; BOLTON & FISHER 2011). This subterranean species tends to nest in sun-exposed sites as well as in woodlands and greenhouses. Both ergatomorphic and winged gynes and males occur.

Occurrence in Iceland: Exclusively found outdoor, with populations confined exclusively to geothermal areas or their immediate vicinity.

*Lasius platythorax* (SEIFERT, 1991) – UI. 2 records

Akranes (1), Garðabær (1)

Eurosiberian species, distributed in mountain zones and also in lowland forests at higher latitudes (SEIFERT 2018). Nests in rotten wood, under moss pads, and in soil under stones, preferentially in deciduous and pine forests. Colonies are monogynous and contain up to 20.000 workers. Strong preference for trophobiosis and is also zoophagous and nectarivorous.

Occurrence in Iceland: One large colony detected in Akranes near a small greenhouse (first noticed in summer 2018); on-site surveys in April 2021 showed hundreds of workers foraging outdoors (likely tending aphids), and in June 2021, tens of gynes and males were observed ready for nuptial flights at nest entrances (homeowners opted against extermination). A second, small colony (one queen, seven workers, and five larvae) was detected in a patch of moss bought in a flower shop.

*Myrmica rubra* (LINNAEUS, 1758) – UI. 1 record Reykjavík (1)

Eurosiberian species, extending North to 70,6°N in Norway. Mainly found in temperate and boreal regions, they nest in urban, agricultural, and natural environments (SEIFERT 2018). Workers excavate rotten wood, litter, and soil under stones. Generally monogynous in early stages but polygynous at maturity, with up to 15 queens and 20.000 workers per nest. Generalist species in terms of diet, but also tend aphids and scale insects.

Occurrence in Iceland: One large colony with two queens in a private garden. First time seen in 2018 or 2019. Successfully eradicated (MANCINI & al. 2022).

### Species known solely from indoor populations

*Monomorium pharaonis* (LINNAEUS, 1758) – IINH & UI. 41 records

Eskifjörður (1, IINH), Garðabær (2, IINH), Grindavík (1, IINH), Grundarfjörður (1, IINH), Hafnarfjörður (1, UI), Keflavík (1, UI), Kópavogur (1 IINH; 1, UI), Reykjavík (23, IINH; 5, UI), Seltjarnarnes (1, IINH), Stykkishólmur (1, IINH), Vogar (1, IINH; 1, UI)

Originally from tropical Asia, it is among the most widespread house ant and can occur in continuously heated buildings (WETTERER 2010b). Typically, does not excavate nest chambers, but instead uses preformed microspaces (e.g., wall, crack, under the floor, behind windowsill) near water sources (KLOTZ & al. 2008). Forms large polygynous colonies with up to 2.000 queens and several hundred thousand workers (SEIFERT 2018). Importantly, this species may transmit pathogens to humans, such as *Salmonella* spp., *Staphylococcus* spp., and *Streptococcus* spp. A highly omnivorous species, actively preying on insects.

Occurrence in Iceland: Exclusively inside continuously heated buildings. Very populous colonies found infesting entire apartment building complexes, mostly near sources of water, in kitchens and bathrooms. We accompanied a pest management professional (SSG) on nine infestations; all nine populations were successfully eradicated.

*Tapinoma melanocephalum* (FABRICIUS, 1793) – IINH & UI. 10 records

Keflavík (2, IINH), Reykjavík (6, IINH & 2, UI)

Widespread tramp species with paleotropical origin (WETTERER 2009) which has spread across the Old and the New World. Considered to be an indoor and outdoor common pest in the tropics and subtropics and common indoor pests in temperate regions. Outdoors, they nest in rotten wood, under bark, and in soil, while indoors, they may nest in cracks and crevices, in potted plants, behind baseboards, and in several other locations where a high level of humidity is guaranteed (KLOTZ & al. 2008). Colonies are highly polygynous, populous, and readily relocate the nest when optimal requirements are not met. Workers are monomorphic and small (<1,5mm). A polyphagous species, with a preference for live and dead small insects and sugary food.

Occurrence in Iceland: Exclusively inside heated buildings. Very populous colonies that can infest entire building complexes, often found in kitchens, living rooms, and laundry rooms. We accompanied a pest management professional (SSG) on two infestations; both populations were successfully eradicated.

*Monomorium* sp. near *M. pharaonis* – UI. 3 records

Hafnarfjörður (2), Reykjavík (1)

Occurrence in Iceland: Large colonies. Often found near sources of water, in kitchens and bathrooms. It can infest entire complexes. To date, all known colonies have been successfully eradicated.

### Species without known established populations

*Camponotus vagus* (SCOPOLI, 1763) – IINH. 8 records

Flateyri (1), Grindavík (1), Kópavogur (1), Reykjavík (2), Selfoss (1), Þorlákshöfn (2)

Western Palearctic species, found from Portugal to Mongolia (SEIFERT 2018). Nests are in rotten wood in woodlands and grasslands. Colonies range from 1.000 to 4.000 individuals, and workers are polymorphic (BENOIS 1972). Zoophagous and aphidicolous. Not commonly found outside its native range.

Occurrence in Iceland: Found in imported construction timber and pallets arrived from abroad. No established colonies or queens found; colony fragments exterminated on-site.

*Camponotus herculeanus* (LINNAEUS, 1758) – IINH. 6 records

Hafnarfjörður (1), Kópavogur (1), Reykjavík (2), Selfoss (1), Þorlákshöfn (1)

Panpalearctic, highly frost-resistant species, common in montane coniferous forests in Central Europe. Nests are often excavated in living or rotten wood. Colonies can be monogynous or oligogynous, populous (up to 40.000 workers), and workers are highly polymorphic (3,5 to 9mm (HEDGES 2010)). This species relies on trophobiosis with aphids for carbohydrate intake. (SEIFERT 2018). It is considered a structural pest (HANSEN & KLOTZ 2005).

Occurrence in Iceland: Found in imported construction timber. Probably queenless colony fragments. No colonies found to date.

*Colobopsis truncata* (SPINOLA, 1808), formerly *Camponotus truncatus* – IINH & UI. 4 records Garðabær (1, UI), Kópavogur (2, IINH), Reykjavík (1, IINH)

Common Mediterranean species, exclusively arboreal, with nesting preference in broad-leaved trees (SEIFERT 2018). Occasionally nesting in construction timber. Colonies are monogynous, polydomous, and small (hardly exceeding 500 workers), and workers are dimorphic (majors have plug-like heads, while minors have normal head morphology). They prey on small insects and arachnids. Weak invaders.

Occurrence in Iceland: Found in wooden goods (e.g., Christmas decorations) and construction timber. Probably queenless colony fragments. No colonies found to date.

*Crematogaster scutellaris* (OLIVIER, 1792) – IINH & UI. 2 records

Akureyri (1, UI), Reykjavík (1, IINH)

Western Mediterranean species, abundantly found in deciduous and conifer forests and gardens and parks near human settlements. Constructs large carton nests in cavities of trees and excavates the bark of cork oaks (SEIFERT 2018). Primarily monogynous, creating large colonies, with a minimum of 20.000 workers. Largely omnivorous.

Occurrence in Iceland: Possible anthropogenous introduction with plant material.

*Lasius emarginatus* (OLIVIER, 1792) – IINH. 2 records

Hafnarfjörður (1), Reykjavík (1)

West Palearctic species, prevalently found in sun-exposed grassy areas (SEIFERT 2018). Nests are mainly found under rocks and in rotten logs, with a cartoon structure in the core of the nest. Strictly monogynous with populous colonies (up to 20.000 workers per nest). This species intensively tends to aphids for honeydew but is also zoophagous and nectarivorous.

Occurrence in Iceland: Total of 2 records in imported goods/plant material; no established permanent colonies.

*Tetramorium caespitum* (LINNAEUS, 1758) – IINH. 2 records

Reykjavík (2)

European species, mainly nests in soil and under stones in grasslands, meadows, riverbanks, and light pine forests (SEIFERT 2018). Colonies are monogynous, with ∼ 15.000 workers, and tend aphids for honeydew, as well as preying on small arthropods and scavenging.

Occurrence in Iceland: Possible anthropogenous introduction with plant material.

*Leptothorax acervorum* (FABRICIUS, 1793) – IINH. 1 record

Reykjavík (1)

Holarctic species, highly tolerant to cold and found in Europe up to 71°N and in the Alps (up to 3.100m). Nests are located under stones and in rotten wood in rocky areas or light forests of *Pinus* (SEIFERT 2018). Colonies are often polygynous, but monogynous is also common. Colonies are small (100-300 workers), and ants actively predate and scavenge for microarthropods (e.g., Collembola).

Occurrence in Iceland: Possible anthropogenous introduction with plant material.

*Monomorium floricola* (JERDON, 1851) – UI. 1 record

Reykjavík (1)

Probably originated in tropical Asia (DEYRUP 2020), is a widespread tramp species (WETTERER 2010a). It occurs in grasslands and agricultural areas, as well as in the urban environment. Highly polygynous large colonies that spread by budding. They prey on small insects and also feed on honeydew. It is not uncommon to see workers in flowers to collect nectar.

Occurrence in Iceland: Single record inside a heated building. Possible anthropogenous introduction with plant material.

*Nylanderia bourbonica* (FOREL, 1886) – UI. 1 record

Kópavogur (1)

Uncertain origin, probably from tropical Australasia (DEYRUP 2020). It is now a widespread tramp species. Extensively found in wet and highly disturbed areas, where it tends to nest on the ground. Colonies are polygynous, and workers actively prey and scavenge small arthropods and tend aphids for honeydew secretion.

Occurrence in Iceland: Single record inside a heated building. Possible anthropogenous introduction with plant material.

*Nylanderia jaegerskioeldi* (MAYR, 1904) – UI. 1 record

Kópavogur (1)

Afrotropical species also extend to Palearctic regions. (LAPOLLA & al. 2011). Outside its original range, it is found in leaf litter and gardens where humidity levels and temperatures are high (GOMEZ 2017; SHARAF & al. 2018).

Occurrence in Iceland: Anthropogenous introduction with plant material.

*Pheidole bilimeki* (Mayr, 1870) – UI. 1 record

Reykjavík (1)

Neotropical species, primarily found in lowland forests and montane areas (LONGINO & COX, 2009). Nests in rotted wood and under stones. Also common on roadsides in the urban environment. Workers are highly polymorphic, with big-headed majors. Generalist diet.

Occurrence in Iceland: Possible anthropogenous introduction with plant material.

*Tetramorium bicarinatum* (NYLANDER, 1846) – UI. 1 record

Garðabær (1)

Uncertain origin, probably from Southeast Asia (BOLTON 1977). A worldwide tramp species, widely distributed by human commerce. They nest under bark, in rotten wood, and in soil (DEYRUP 2020). Colonies are highly polygynous, and they spread by budding. Colonies are populous. They eat living or dead insects and tend to sap-sucking insects for honeydew.

Occurrence in Iceland: Anthropogenous introduction with plant material.

*Trichomyrmex destructor* (JERDON, 1851), formerly *Monomorium destructor* – IINH. 1 record Seltjarnarnes (1)

Presumably native to the Indian subcontinent (DEYRUP 2020). It is a widespread tramp species distributed in tropical and subtropical regions. Nesting preference is shown for disturbed areas, in soil, trees, and inside buildings. Colonies are highly polygynous and workers are polymorphic. Generalist species.

Occurrence in Iceland: Anthropogenous introduction with wooden goods.

*Camponotus* sp. – IINH. 1 record

Reykjavík (1)

Occurrence in Iceland: Queenless colony fragment, apparently arrived with goods shipped from India.

*Pheidole* sp. – IINH. 1 record

Reykjavík (1)

Occurrence in Iceland: Anthropogenous introduction with plant material.

Myrmicinae (subfamily), species a – IINH. 1 record

Reykjavík (1)

Myrmicinae (subfamily), species b – IINH. 1 record

Kópavogur (1)

### Known ant distribution in Iceland

Most ant records in Iceland were from the Capital Region (575; 88.60% of the total), with the geographically neighboring regions having several cases (26 and 19 in Southern Region and Southern Peninsula, respectively) (Fig. 2 & Tab. 2). The Eastern Region and the Westfjords had very few cases (1 and 2, respectively). In 95.69% of the cases (621 records), ants were detected indoors in continuously heated buildings. Consistent with predictions of ÓLAFSSON & RICHTER (1985), the Húsamaur (“house ant” in Icelandic) species *H. ergatandria* (previously misidentified as *H. punctatissima*) was most common (516 records; 79.51% of the total; Tab. 2). *H. ergatandria* was almost exclusively found in urban habitats, and had densest distribution around Reykjavík, reaching east to Vík í Mýrdal and north to Reykholt in Borgarfjörður. There was also a concentration of this species in Akureyri, the largest town in north of Iceland, and villages nearby. No cases have been reported from the Westfjords, or regions east of Akureyri and Vík í Mýrdal.

**Fig. 2:**
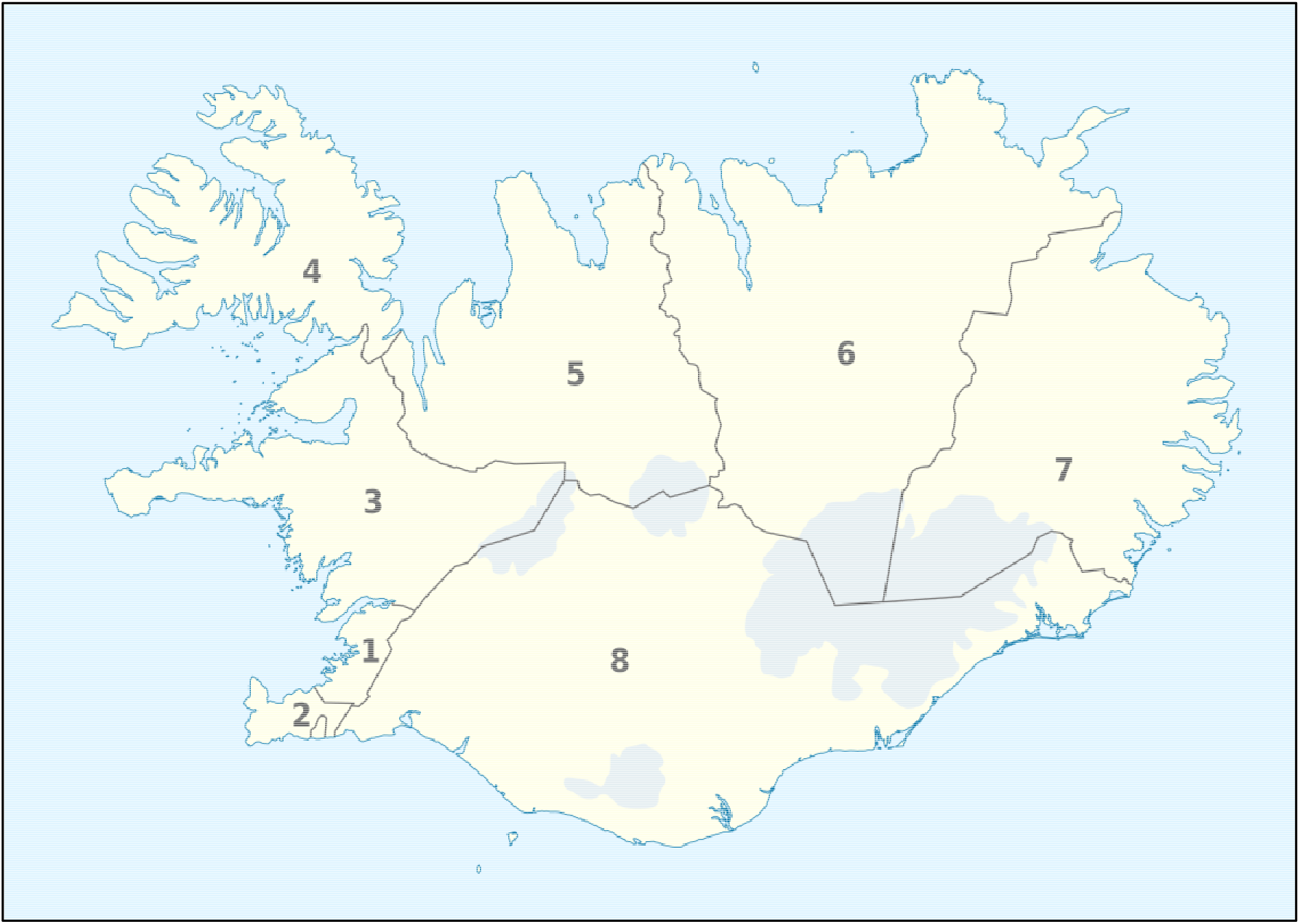
Map of Iceland divided into administrative regions. The regions are numbered as follows: 1. Capital Region, 2. Southern Peninsula, 3. Western Region, 4. Westfjords, 5. Northwestern Region, 6. Northeastern Region, 7. Eastern Region, and 8. Southern Region. Source: TUBS, adapted from Wikipedia (https://en.wikipedia.org/wiki/Regions_of_Iceland) licensed under CC BY-SA 3.0.

**Fig. 3:**
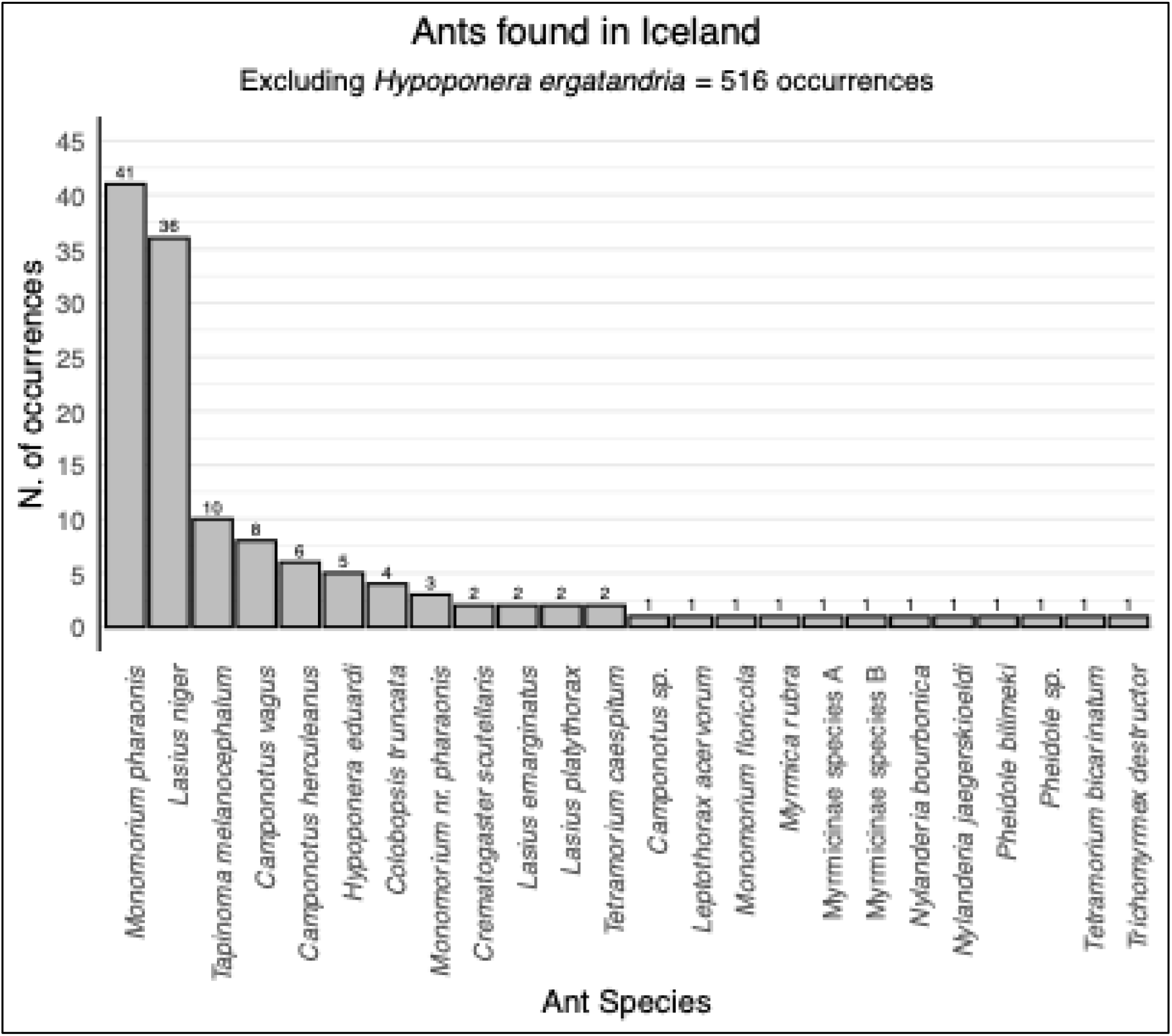
Frequency distribution of ant records in Iceland between 1974 and 2022, based on combined data from the Icelandic Institute of Natural History (IINH) and University of Iceland (UI) collections. The x-axis lists the ant species identified, while the y-axis shows the number of records for each species. Species are ordered in descending frequency, from left to right. Note that *Hypoponera ergatandria* is excluded from this figure, as its 516 occurrences would significantly exceed the scale and distort the visualization.

**Tab. 2:** Ant records organized by species (rows) and geographical regions within Iceland (columns). Both ant species and geographical regions are arranged in descending order of record counts. The total counts and percentages of all records (combining IINH and UI data) are provided for each species (on the right) and region (at the bottom).

| N. | Ant species | Capital Region | Southern Region | Southern Peninsula | Western Region | Northeastern Region | Northwestern Region | Westfjords | Eastern Region | Records per species | Percent of the total (%) |
|---|---|---|---|---|---|---|---|---|---|---|---|
| 1 | *Hypoponera ergatandria* | 473 | 13 | 10 | 8 | 6 | 5 | 1 | - | **516** | 79.51 |
| 2 | *Monomorium pharaonis* | 34 | - | 4 | 2 | - | - | - | 1 | **41** | 6.32 |
| 3 | *Lasius niger* | 28 | 3 | 2 | 3 | - | - | - | - | **36** | 5.55 |
| 4 | *Tapinoma melanocephalum* | 8 | - | 2 | - | - | - | - | - | **10** | 1.54 |
| 5 | *Camponotus vagus* | 3 | 3 | 1 | - | - | - | 1 | - | **8** | 1.23 |
| 6 | *Camponotus herculeanus* | 4 | 2 | - | - | - | - | - | - | **6** | 0.92 |
| 7 | *Hypoponera eduardi* | - | 5 | - | - | - | - | - | - | **5** | 0.77 |
| 8 | *Colobopsis truncata* | 4 | - | - | - | - | - | - | - | **4** | 0.62 |
| 9 | *Monomorium* nr. *pharaonis* | 3 | - | - | - | - | - | - | - | **3** | 0.46 |
| 10 | *Crematogaster scutellaris* | 1 | - | - | - | 1 | - | - | - | **2** | 0.31 |
| 11 | *Lasius emarginatus* | 2 | - | - | - | - | - | - | - | **2** | 0.31 |
| 12 | *Lasius platythorax* | 1 | - | - | 1 | - | - | - | - | **2** | 0.31 |
| 13 | *Tetramorium caespitum* | 2 | - | - | - | - | - | - | - | **2** | 0.31 |
| 14 | *Camponotus* sp. | 1 | - | - | - | - | - | - | - | **1** | 0.15 |
| 15 | *Leptothorax acervorum* | 1 | - | - | - | - | - | - | - | **1** | 0.15 |
| 16 | *Monomorium floricola* | 1 | - | - | - | - | - | - | - | **1** | 0.15 |
| 17 | *Myrmica rubra* | 1 | - | - | - | - | - | - | - | **1** | 0.15 |
| 18 | *Nylanderia bourbonica* | 1 | - | - | - | - | - | - | - | **1** | 0.15 |
| 19 | *Nylanderia jaegerskioeldi* | 1 | - | - | - | - | - | - | - | **1** | 0.15 |
| 20 | *Pheidole bilimeki* | 1 | - | - | - | - | - | - | - | **1** | 0.15 |
| 21 | *Pheidole* sp. | 1 | - | - | - | - | - | - | - | **1** | 0.15 |
| 22 | *Tetramorium bicarinatum* | 1 | - | - | - | - | - | - | - | **1** | 0.15 |
| 23 | *Trichomyrmex destructor* | 1 | - | - | - | - | - | - | - | **1** | 0.15 |
| 24 | Myrmicinae, species a | 1 | - | - | - | - | - | - | - | **1** | 0.15 |
| 25 | Myrmicinae, species b | 1 | - | - | - | - | - | - | - | **1** | 0.15 |
| **Records per region** |  | **575** | **26** | **19** | **14** | **7** | **5** | **2** | **1** | **649** |  |
| Percent of the total (%) |  | 88.60 | 4.01 | 2.93 | 2.16 | 1.08 | 0.77 | 0.31 | 0.15 |  |  |

While *H. ergatandria* was almost exclusively found indoor in urban habitats, *H. eduardi* and *L. niger* were more common outdoors (Tab. 3). All but 28 records, representing 5 species, were exclusively found indoors. The exceptions to this are *L. niger* (12/36 in gardens), *H. eduardi* (4/5 in geothermal areas and 1/5 in garden), *H. ergatandria* (8/516 in geothermal areas, and 1/516 in an urban garden), *L. platythorax* (1/2 in an urban garden) and *M. rubra* (1/1 in an urban garden).

**Tab. 3:** Ant records in Iceland collected between 1974 and 2022 (IINH and UI collections), categorized by habitat (indoor or outdoor). Ant species are listed in descending order based on the number of records, with total counts per species also provided.

| **N.** | **Ant species** | **Habitat** |  | **Total records** |
|---|---|---|---|---|
|  |  | Indoor | Outdoor |  |
| 1 | *Hypoponera ergatandria* | 507 | 9 | 516 |
| 2 | *Monomorium pharaonis* | 41 | - | 41 |
| 3 | *Lasius niger* | 24 | 12 | 36 |
| 4 | *Tapinoma melanocephalum* | 10 | - | 10 |
| 5 | *Camponotus vagus* | 8 | - | 8 |
| 6 | *Camponotus herculeanus* | 6 | - | 6 |
| 7 | *Hypoponera eduardi* | - | 5 | 5 |
| 8 | *Colobopsis truncata* | 4 | - | 4 |
| 9 | *Monomorium* nr. *pharaonis* | 3 | - | 3 |
| 10 | *Crematogaster scutellaris* | 2 | - | 2 |
| 11 | *Lasius emarginatus* | 2 | - | 2 |
| 12 | *Lasius platythorax* | 1 | 1 | 2 |
| 13 | *Tetramorium caespitum* | 2 | - | 2 |
| 14 | *Camponotus* sp. | 1 | - | 1 |
| 15 | *Leptothorax acervorum* | 1 | - | 1 |
| 16 | *Monomorium floricola* | 1 | - | 1 |
| 17 | *Myrmica rubra* | - | 1 | 1 |
| 18 | *Nylanderia bourbonica* | 1 | - | 1 |
| 19 | *Nylanderia jaegerskioeldi* | 1 | - | 1 |
| 20 | *Pheidole bilimeki* | 1 | - | 1 |
| 21 | *Pheidole* sp. | 1 | - | 1 |
| 22 | *Tetramorium bicarinatum* | 1 | - | 1 |
| 23 | *Trichomyrmex destructor* | 1 | - | 1 |
| 24 | Myrmicinae, species a | 1 | - | 1 |
| 25 | Myrmicinae, species b | 1 | - | 1 |

## Discussion

Six alien ant species were recently found in the Faroe Islands (HAMMER & JENSEN 2021): *L. niger*, *M. pharaonis*, *T. melanocephalum*, *Camponotus fallax*, *Formica rufa* and *Tapinoma ibericum*, the first three of which were also found in Iceland. For its peculiarities and the remoteness of the location, the Icelandic biome is vulnerable to the introduction of non-native species, and even the settlement of a single species may have dramatic effects on the island’s biodiversity (SCHMALENSEE 2010; WASOWICZ & al. 2013).

There are two potential explanations for the high concentration of ants in the Capital Region: (a) The Capital Region hosts ∼60% of the total human population in an area of ∼1.000 km^2^ with connected towns (also the main port of entry for most of the imported goods and plant material); (b) In other regions, ant diversity and distribution may be underestimated since pest management professionals and individual citizens may be unaware of ants and/or less willing to report ant cases to IINH or UI. All of them can be considered alien or causal alien to Iceland, with possible exception of *H. eduardi*. Two species may be considered naturalized.

WETTERER (2015) summarized 42 cosmopolitan ant species, including the ones that already had a cosmopolitan spread at the beginning of the 20^th^ century (FOREL 1911) and others that spread more recently. Eight of the 25 species in Iceland are on that list: *Hypoponera eduardi*, FOREL, 1894; *Monomorium floricola*, JERDON, 1851; *Monomorium pharaonis*, LINNAEUS, 1758; *Myrmica rubra*, LINNAEUS, 1758; *Nylanderia bourbonica*, FOREL, 1886; *Tapinoma melanocephalum*, FABRICIUS, 1793; *Tetramorium bicarinatum*, NYLANDER, 1846; *Trichomyrmex destructor*, JERDON, 1851. These are defined as highly invasive ant species, characterized by five typical traits that have led to the term tramp species (HÖLLDOBLER & WILSON 1990). Five of these species are defined as casual aliens in Iceland (PYŠEK & al. 2004) and we believe they do not currently have living populations on the island. While four of those species (*M. floricola*, *N. bourbonica*, *T. bicarinatum*, and *T. destructor*) were exclusively detected inside heated buildings and would have no chance of surviving outdoors in Iceland, the *M. rubra* colony constitutes a special case since a large colony with two queens was found in a garden, and the foraging activity was seemingly performed entirely outdoors (MANCINI & al. 2022).

The remaining three species of cosmopolitan ants, deserve particular attention. *M. pharaonis* and *T. melanocephalum* were already included in FOREL’s list (1911) and are now global pests. In Iceland, they have been found infesting entire apartment buildings. Following the classification provided by PYŠEK & al. (2004), these two species represent casual alien species since single populations may occasionally spread, but the species need repeated introductions to persist in the invaded environment.

The data on the third species, *H. eduardi*, show that their populations are exclusively confined to geothermal areas, or the near vicinity. Thus, at this stage, we cannot conclude if this species has arrived in Iceland through human-assisted pathways or, alternatively, if it naturally colonized the island before human settlement. We consider *H. eduardi* to be a cryptogenic species (ESSL & al. 2018) until further evidence is gathered.

Given its cryptic behavior, a remarkably complicated species to eradicate is *H. ergatandria*. Regarding the biology of the species, it is very challenging to identify the location of *H. ergatandria* nests due to the cryptic habits of the workers, and even when ants are detected jumping/flying toward the windowsills attracted by the light, these are only newly emerged winged gynes. Ant gynes do not usually feed while searching for a new location to establish their nests; therefore, baiting stations are not a suitable pest management solution. The only possible way to eradicate *H. ergatandria* nests is by breaking up the floors and inserting protein food coated with avermectin dust (KLOTZ & al. 2008, referring to *H. punctatissima*). However, this would represent a costly procedure for the homeowners and does not guarantee extermination success because of the species’ habit of creating “sister nests” a few meters from the original nest, making it almost impossible to identify all the locations of functioning colonies (or sub-colonies). Therefore, we expect more *H. ergatandria* cases to be reported in the future. However, the question remains if the public and pest management professionals become less inclined to report the cases. *H. ergatandria* may be considered naturalized alien species in Iceland since it reproduced and self-sustained populations for more than ten years (PYŠEK & al. 2004).

Among the *Lasius* spp. found, *L. platythorax* can be considered a casual alien species (PYŠEK & al. 2004) in Iceland. *L. niger* is a common species in temperate Europe, and it seems it may be able to adapt to the Icelandic “garden”-environment successfully. It is still unclear if this species may be considered naturalized (PYŠEK & al. 2004) in Iceland, and more data are needed.

Carpenter ants (*Camponotus* spp.) and several other rarely detected ant species in Iceland, such as *L. emarginatus* and *P. bilimeki*, appear to represent only occasional introductions via imported goods or timber, with no evidence of established permanent colonies. Consequently, both the carpenter ants and these rare species can reasonably be considered casual alien species (PYŠEK & al. 2004).

The genetic data generally reveal limited genetic diversity within Iceland, suggesting a single colonization event for each of the ant species detected. It may be possible to generate a hypothesis about their geographic origin if a sufficient collection of records is available from other countries such as in the case of *M. rubra* found in Iceland (MANCINI & al. 2022). The highest within-species variation was in *L. niger* records, which may correlate with outdoor vs indoor colonies or repeated colonization (though the sample size is very limited). As an anecdote, we like to bring attention to the deep branch among *Monomorium* spp. This level of sequence divergence is normally associated with speciation. The fact that both branches were found in such a remote place as Iceland suggests that two *Monomorium* species may be cosmopolitan tramps and urges further study of the genetic diversity of this genus.

### Conclusions and future research

Because of the location of Iceland in the middle of the North Atlantic, the geological origin of the landmass, and, importantly, the repeated ice-age glaciations, none of the 25 ant species found on the island can be considered native at this stage. In accordance with ESSL & al. (2018), all species, but one, can be considered alien or casual alien to the island since (a) we know that biogeographical boundaries that, in the case of an oceanic island, correspond to administrative boundaries, have been crossed, (b) there is suggestive evidence of direct human agency in the transport of the species, (c) these species survive without human assistance, and (d) we collected information on the year of the first finding for every species and we can, therefore, distinguish species as alien or casual alien. One species, *H. eduardi*, may be considered cryptogenic (CARLTON 1996; ESSL & al. 2018), since there is currently no direct evidence for it to be considered alien and further investigation is needed. One of the alien species (*H. ergatandria*) can be considered naturalized in Iceland. Analysis of the genetic constitution and relatedness of populations around the island may in the future establish if they reflect one or more colonizations, and patterns of migration. The majority of the detected ant species (a) most likely colonized the island in the last half-century, and (b) seem to be strictly linked to heated buildings in the urban environment. Developing and implementing more extensive public outreach programs and expanding and strengthening relationships with pest management professionals will give access to ever-increasing numbers of ant cases in Iceland and improve our knowledge of ant diversity and distribution. Future focus of this work could be to (a) look at pathways of invasions and ways to manage them, (b) develop monitoring strategies to help detect new invasions and track old ones, (c) assess potential impacts of these ant species on native biomes, and (d) develop better biosecurity strategies.

## Acknowledgments

We thank Erling Ólafsson and Matthías Svavar Alfreðsson, for collecting, cataloging, and identifying ant findings and giving us access to the Icelandic Institute of Natural History (IINH) dataset and consultation on ant matters; the pest management professional Steinar Smári Guðbergsson, for allowing us to join him during his operations; Bernhard Seifert, for identifying most of the specimens, John T. Longino for identifying *Pheidole* sp. specimens and John LaPolla for *Nylanderia* spp.; and Mattia Menchetti for providing unpublished comparative sequence data; and several individual citizens, for assisting us in collecting ants.

## Authors’ contributions

MM conceived and planned the project and wrote the manuscript draft. MM and AGG collected data. MM, JKW, and AP contributed to project planning. MM and AP analyzed the data. AP, and JKW, revised manuscript. All authors approved the submitted version and agreed to be held accountable for all aspects of the work.

## Declaration of generative AI usage

The authors declare that no generative artificial intelligence tools were used at any stage of preparing or writing this manuscript.

